# Ghat: An R package for identifying adaptive polygenic traits

**DOI:** 10.1101/2020.07.06.189225

**Authors:** Medhat Mahmoud, Mila Tost, Ngoc-Thuy Ha, Henner Simianer, Timothy Beissinger

## Abstract

Identifying selection on polygenic complex traits in crops and livestock is important for understanding evolution and helps prioritize important characteristics for breeding. The QTL that contribute to polygenic trait variation often exhibit small or infinitesimal effects. This hinders the ability to detect QTL controlling polygenic traits because enormously high statistical power is needed for their detection. Recently, we circumvented this challenge by introducing a method to identify selection on complex traits by evaluating the relationship between genome-wide changes in allele frequency and estimates of effect-size. The approach involves calculating a composite-statistic across all markers that captures this relationship, followed by implementing a linkage disequilibrium-aware permutation test to evaluate if the observed pattern differs from that expected due to drift during evolution and population stratification. In this manuscript, we describe “Ghat”, an R package developed to implement this method to test for selection on polygenic traits. We demonstrate the package by applying it to test for polygenic selection on 15 published European wheat traits including yield, biomass, quality, morphological characteristics, and disease resistance traits. Moreover, we applied Ghat to different simulated populations with different breeding history and genetic architecture. The results highlight the power of Ghat to identify selection on complex traits. The Ghat package is accessible on CRAN, the Comprehensive R Archival Network, and on GitHub.

## Introduction

Many important traits in plants, animals, and humans are polygenic and depend on the cumulative effects of many loci, each contributing a small proportion of the total genetic variation [1]. Studying the evolution and selection history of such traits can be challenging due to the small effects of the quantitative trait loci (QTL) that contribute to genetic variation. However, regardless of effect size, the theory of natural selection tells us that alleles with positive effects on fitness will tend to increase in frequency over successive generations [2]. Several statistical methods have been developed to identify loci carrying such beneficial alleles. These methods were reviewed in [3], and include approaches relying on polymorphism variability, haplotype frequency, linkage disequilibrium, allele frequency change over time, and techniques that combine some or all of these signals. However, in the case of highly polygenic traits, it is difficult or impossible to identify regions carrying individually selected alleles/haplotypes due to their small effects, regardless of which sophisticated methodologies are leveraged. For instance, it has been shown that selected loci are difficult to identify when per-locus selection intensity is low [3], as is the case with low heritability traits and those controlled by many alleles of small effect [4]. This imposes serious limitations to the previous methods when identifying selection on polygenic traits (e.g. [5]).

Recently, large sample sizes, especially as utilized in human genetic studies, have allowed a more powerful understanding of the genetic architecture of polygenic traits using genome-wide association studies (GWAS) [6–8]. After conceptual work solidified the importance of polygenic adaptation [9,10], researchers began placing more of an emphasis on identifying this phenomenon using an infinitesimal approach [11–14]. When polygenic adaptation is studied with an infinitesimal approach, the estimates of individual allelic effects on a phenotype are often calculated in a GWAS[12,15,16]. The “GWAS hits” are then used to test for selection [12,15,16]Studies in human genetics usually use summary statistics from meta-analysis GWAS like the GIANT data set, where many individual GWAS are combined [12,15–19]. These studies observed strong signals for polygenic selection [12,15–19]]. The observed signals did not replicate when the estimates of individual allelic effects were calculated based on GWAS conducted in the unstructured UK Biobank data set, especially when the data set was filtered for individuals with a common ancestry [15,16]. Apparently estimates of allelic effects derived from GWAS are affected by population confounding effects, and when they are combined in polygenic scores, they are highly prone to cumulative bias due to their systematic nature [15,16,19]. Furthermore, Berg et al. (2019) [16] observed that methods like principal component analysis are not properly accounting for population confounding effects.

Zeng et al. (2018) [20] recently developed a Bayesian mixed linear model (BayesS) based on the relationship between SNP effect sizes and minor allele frequency. This approach uses genomic data and fits all SNP effects together as random effects [20]. The identified signals of polygenic selection from Zeng et al. (2018) [20] conflict with the results from Berg et al. (2019) [16], even though both studies worked with the unstructured UK Biobank data set. Berg et al. (2019) [16] filtered the samples based on British ancestry, whereas Zeng et al. (2018) [20] included samples of all European ancestry, which might suggest that residual European population structure continues to be confound, even when advanced methods for corrections for population structure were applied [16]. On the other hand, BayesS used by Zeng et al. (2018) [20] is based on the relationship between SNP effect sizes and minor allele frequency, whereas minor allele frequency and SNP effects might correlate due to estimation error during genomic prediction [4].

The previously mentioned approaches were developed for human genetics, and most of these approaches are based on summary statistics from massive GWAS which are not always feasible for other species, e.g. due to budgetary limitations. Josephs et al. (2019) [21] developed a statistic, referred as conditional Q_PC_, which detects adaptive divergence while controlling for population substructure. Conditional Q_PC_ detects adaptive divergence based on the excess of variation observed among populations versus variation observed within populations [21]. The underlying assumption of this test is that adaptive divergence among populations increases the trait variance explained by the first principal components (PCs) relative to the later PCs [21]. The PCs are computed based on the relatedness matrix, therefore the conditional Q_PC_ detects substructures which are associated with trait divergence and based on these observations, conclusions about polygenic adaptation regarding the trait are drawn [21].

Another test, denoted *Ĝ* (or Ghat), has been developed to identify selection on polygenic traits using whole-genome data [4]. Ghat combines information from all loci simultaneously to evaluate whether there is a trend for loci contributing to a phenotype to collectively show evidence of selection [4,22]. Thereby, signals that are individually insignificant for selection (at the locus level) may become highly significant when evaluated at the genome-wide scale (at the level of finding selection on a trait). Furthermore, because Ghat uses pre- and post-selection data, it can identify polygenic selection within shorter sub-periods of time and compare these to each other [4]. Another advantage of Ghat is that it is based on the relationship between SNP effects and allele frequency change over time at every locus, which should not correlate unless there has been on-going selection in comparison to minor allele frequency and SNP effects [4,20].

We have developed and released an R [22,23] package called “Ghat” to implement the *Ĝ* test. The package allows users to test for selection and evolution in quantitative traits, and as shown by Beissinger et al. (2018) [4], it is particularly powerful when testing for selection on highly polygenic traits. To evaluate and demonstrate the package in this manuscript, we applied the method and corresponding Ghat package to 15 different traits measured in a winter wheat collection of 191 cultivars, registered mainly in Western Europe between 1966 and 2013, which were published by Voss-Fels et al. (2019) [24]. We then applied Ghat package on a simulated cattle population to validate the package and explore different situations where it is most and least powerful.

## Implementation of the Ghat Package

### Overview

Ghat identifies selection by leveraging all genotyped loci simultaneously [4]. This makes it possible to identify selection in complex traits that are controlled by many genes with small effects [4]. First, all allelic effects are estimated using a genomic prediction approach. Next, allele frequencies are estimated in two or more different generations (e.g. generation 0 and 10) [4]. Third, the rate of linkage disequilibrium decay is estimated to approximate the number of independent genome segments in the study population [4]. Finally, an LD-aware permutation test is implemented to evaluate if there is evidence of significant selection between generations. The direction of selection is calculated based on the relationship between the change in allele frequency between the generations and the estimated additive effect [4]. Complete theoretical and methodological details are provided in [4].

### Installation

The Ghat package is available on the Comprehensive R Archival Network (CRAN) [23] and can be installed within the R terminal (Fig1, step-1). The package requests the following optional R-package dependencies:

- Parallel-Package {parallel}: Support for parallel computation, to speed the computation time in large data analysis. [25]
- rrBLUP-Package {rrBLUP}: Ridge regression for estimating marker effects (RR-BLUP). [26]

**Fig 1.**
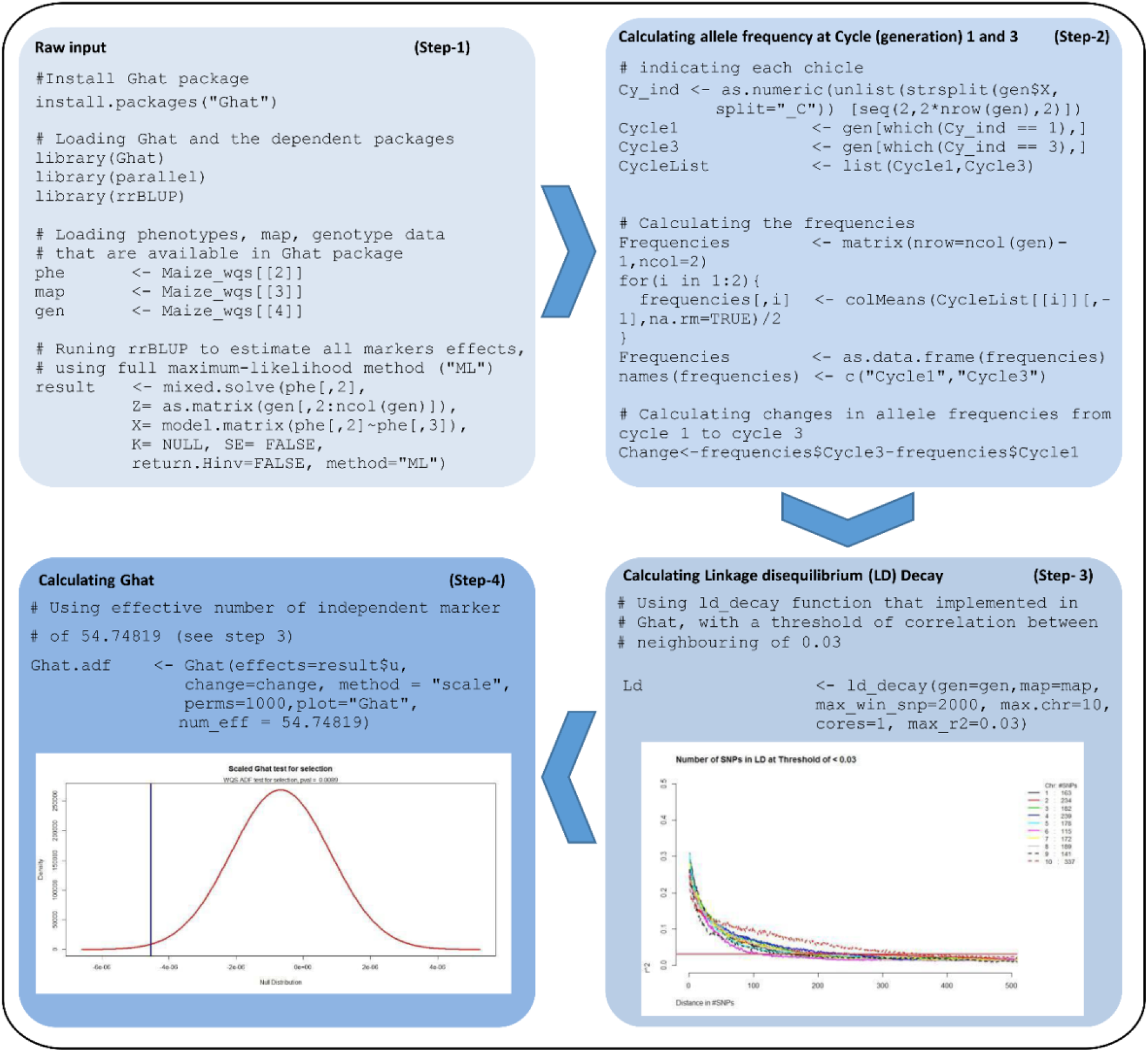
Four steps to go from raw individual marker data to results from the Ghat test for selection. An illustrated example using the “Maize_wqs” data available in the Ghat package. **Step-1**: install the package, load the dependencies and “Maize_wqs” data and estimate allele substitution effects (e.g., by using rrBLUP [26]). **Step-2**: calculate the differences in allele frequency between two different generations (cycle 1 and cycle 3). **Step-3**: estimate the decay of linkage disequilibrium (LD) to calculate the effective number of independent markers. **Step-4**: calculate the Ghat value and p-value from the permutation test.

These maybe installed and loaded before running Ghat by typing:

>install.packages(c(“parallel", “rrBLUP”)) and then typing >library(parallel) and >library(rrBLUP) (Fig1, step-1).

### Input Data format

For its simplest implementation, Ghat requires only two vectors of input. The first of these is a vector of allele substitution effects, and the second is a vector containing the allele frequency change between the two generations (Table 1) [4]. Required parameters are, “method”: to specify how the Ghat permutation test should account for LD between markers (“scale” is usually the most appropriate method to implement because it accounts for linkage disequilibrium between markers, as described below), “perms”: for the number of permutations to perform, “plot”: to specify whether or not simple Ghat plots should be output, “blocksize”: for setting the size of blocks that should be used for LD-trimming (only required when method = “trim”), and “num_eff”: for setting the effective number of independent segments of the genome (only required when method = “scale”). All parameter details are described in Table 1.

**Table 1.** Objects, data, functions, and options found in Ghat package.

| Function |  | Description |
| --- | --- | --- |
| Ghat<br>(Function) | effects | Input vector of allele substitutional effects. |
|  | change | Input vector of changes in allele frequency between two different generations (could be positive, negative or zero). |
|  | method | One of "vanilla" (assumes complete linkage equilibrium between markers), "trim" (excludes markers to approximate linkage equilibrium some of the extreme values), or "scale" (scales results to reflect underlying levels of linkage LD). |
|  | perms | Input number to set the number of permutations to test for significance. |
|  | plot | Input option to select which plot you need to be returned: "Ghat", "Cor", or "Both". |
|  | blockSize | Input number to set the size of the window on each chromosome for trimming, only required if method = "trim". |
|  | num_eff | Input number to set the effective number of independent markers, only required if method = "scale". Can be calculated using "ld_decay" function above. |
| ld_decay<br>(Function) | gen | Input matrix of genotype data. Individuals in rows, genotypes (0, 1, 2) in columns. |
|  | map | Input data frame including the name for each marker with a corresponding chromosome number and physical position. |
|  | max_win_snp | Input number to set the maximum number of markers allowed per window within a chromosome before estimating the LD. The default value is 2000. |
|  | max.chr | Input number to select to which chromosome you want to run the analysis, chromosomes above this number will be excluded from the analysis. |
|  | cores | Input number to set the optimal number of cores for using parallelized calculation, since calculation of LD require too much memory. The default value is 1. |
| | max_r2 | Input number to set the threshold of correlation between the neighbourhood variant ($r^2$) to calculate the effective number of independent variants. |
| Maize_wqs<br>(Data object) | Maize_wqs[[1]] | Data frame, including all SNP names (SNP Names), effects on Acid detergent fiber (adf) (effects. adf), and changes in allele frequencies (change) between cycle 1 and cycle 3 of selection. |
|  | Maize_wqs[[2]] | Data frame, including plant ID's (taxa), estimated breeding values for Acid detergent fiber (adf), and generation when the phenotyped strains were grown (year). |
|  | Maize_wqs[[3]] | Data frame, map file, each line of the Map file describes a single marker and contain three columns. 1: Name (SNP id); 2: Chromosome (Chromosome number); 3: SNP position (in base-pairs). |
|  | Maize_wqs[[4]] | Data frame, Genotype (Illumina MaizeSNP50 BeadChip); an Infinium HD assay (Illumina, Inc. San Diego, CA). 10,017 SNP markers (0, 1 and 2) after filtration, distributed across the maize genome [27]. |

### Establishing the number of effective markers

Treating each marker as an independent observation can lead to overestimated test statistics. This is because allele frequencies and effect estimates from markers that are in LD are not independent [4]. Any level of LD between markers leads to fewer independent observations than the total number of markers [4]. To avoid such inflated signal of polygenic adaptation, we include an option in Ghat package called “scale”, which scales the variance of the permutation test statistic according to the realized extent of LD [4]. More details on the statistical background of scaling method were published by Beissinger et al. (2018) [4]. In the Ghat package, we provide a function called “ld_decay” that can estimate the LD decay across the entire genome and uses this information to approximate the total number of independent genome segments, or “genotype blocks” [4]. The following command runs the ld_decay function >Ld <-ld_decay(gen, map, max_win_snp, max.chr, cores=1, max_r2) (Fig 1, Step-3)[4]. The output information is used by Ghat to scale the variance of the permutation test statistic according to the actual number of independent markers (Fig 1, Step-4) [4].

### Running the test

To run Ghat, estimates of allele substitution effects and allele frequency changes between two generations (or populations) are required [4]. Next, to perform the Ghat test with this information, only one line of code is required: >Ghat_res <-Ghat(effects, change, method, perms, plot, num_eff) (Fig 1, Step-4) [4]. All of the functions, options and data sets available in Ghat are provided in Table 1

## Part I: Testing for selection during 50 years of wheat breeding

To demonstrate the power and flexibility of the Ghat R package, we used it to test for selection during the past 50 years of wheat breeding in Western Europe. To achieve this, we leveraged a large dataset representing 50 years of commercial winter wheat varieties that was previously published by Voss-Fels and colleagues [24]. Our analysis methods and results are described below.

## Methods

### Genotypes and phenotypes

A data set of 191 cultivars, registered mainly in Western Europe between 1966 and 2013, was previously published by Voss-Fels and his group[24]. In total, 19 traits were considered and used to evaluate and quantify the phenotypic progress of wheat breeding during the last five decades. The cultivars in the data set are mainly registered and bred for the German market [24].

After filtering for data quality, 15 traits remained in the data set. Spikes per sqm, heading, sedimentation, falling number, and green canopy duration were removed from the analysis because of missing phenotypic information (>70% of total phenotypes were missing). All traits were tested in three different environments by Voss-Fels et al. (2019) [24]. The environments were: 1: A high-intensity nitrogen supply along with best-practices fungicide, insecticide, and growth regulator applications (HiN/HiF treatment), 2: A high level of nitrogen fertilization with a fungicide-free treatment (HiN/NoF), and 3: A low level of nitrogen fertilization with a fungicide-free treatment (LoN/NoF). The evaluation was based on five main groups of traits: **1- Yield parameters traits**, including grain yield, harvest index, radiation use efficiency and radiation interception efficiency; **2- Biomass parameter traits**, including above-ground biomass, kernels per sqm, and thousand-kernel weight. **3- Quality parameter traits**, including crude protein, protein yield and nitrogen use efficiency. **4- Morphological traits**, including plant height, green canopy duration and kernel per spike. **5- Disease resistances traits**, including resistance to powdery mildew and resistance to stripe rust. All cultivars were genotyped with a 15K SNP Illumina Infinium iSelect genotyping array [28] by Voss-Fels et al. (2019) [24]. After filtering, 8,710 SNPs passed the genotype quality control criteria [24] and were included in their subsequent analyses. In our analysis, we re-analyzed the exact set of SNPs that was published by Voss-Fels et al. (2019) [24], as described above. All estimates of genetic correlation between the same traits measured in different environments were published [24].

### Phenotypic progress

For establish a baseline for progress that we could use to compare against Ghat results, we re-estimated the phenotypic progress of the 15 wheat traits over 50 years of selection and adaptation using simple *Pearson* correlation between phenotype and the year of release [29].

### Ghat

Allelic effects at every marker locus for the 15 traits were estimated using rrBLUP [26], and changes in allele frequencies were calculated with R [23,30]. Ghat is calculated as the summation of the estimated allele frequency change of every SNP multiplied by its effect size, according to

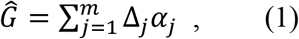

where Δ_*j*_ is the frequency change in locus *j* between two time-points and *α*_*j*_ is the additive effect of locus *α*. We implemented two different analyses with Ghat to test for selection, which enabled us to measure the efficiency of the test in different situations. In the first analysis, which we call “**All Phenotypes”**, allelic effects were estimated using phenotypic information from all available cultivars in the Voss-Fels et al. (2019)[24] study, which represent the last 50 years of wheat breeding. This represents the best-case Ghat analysis, in which all phenotypic information is available. In the second analysis, which we call **“Modern Phenotypes”**, allelic effects were estimated using only modern phenotypic information (from 2010 to 2013), while changes in allele frequency were still calculated based on genotypes from opposite ends of the 50-year dataset. This represents a realistic use-case for Ghat, when all genotypic information is available, but phenotypic information is only available for a subset of the individuals (e.g. from modern germplasm). In other words, this mimics the situation where reliably measured historical phenotypes do not exist, but historical tissues or seeds are available and can be used for DNA collection.

### PCA

Principal component analysis (PCA) was performed with an R [23,31] package called “adegenet”, which was developed for multivariate analysis of genetic markers [31]. The PCA was conducted with all 191 cultivars with 8,710 markers to detect the presence of population structure. For this analysis missing data were imputed according to the mean allele frequency [32].

## Results

### Testing for selection during 50 years of wheat breeding

The main theory behind this test for polygenic selection is that alleles at every locus, distributed across the entire genome, may contribute a small proportion of the total variance for each trait. Therefore, previous studies either required large sample sizes [5] or were unsuccessful in identifying the signals of selection on a genetic level [5]. To demonstrate the Ghat package, we used it to test for selection on the 15 wheat traits, enabling us to establish how much power is lost when phenotypic data are only available from modern lines.

#### Phenotypic progress

After estimating the phenotypic trend (correlation between phenotypic value and the registration year) (Fig 2). We can classify the 15 traits into three groups. The first group is showing comprised of traits that increased in value during breeding. These were above-ground plant biomass, grain yield, green canopy duration, harvest index, kernels per spike, kernels per square meter, nitrogen use efficiency, protein yield, radiation interception efficiency, radiation use efficiency, resistance to powdery mildew, and resistance to stripe rust. Next, we identified a group of traits with a decreasing phenotypic trend, which was comprised of plant height and crude protein. The last group contained traits without a strong overarching phenotypic trend. Only thousand-kernel weight was in this group. The correlation for thousand-kernel weight was positive, but relatively weak in comparison correlations observed in the first group (Fig 2).

**Fig 2.**
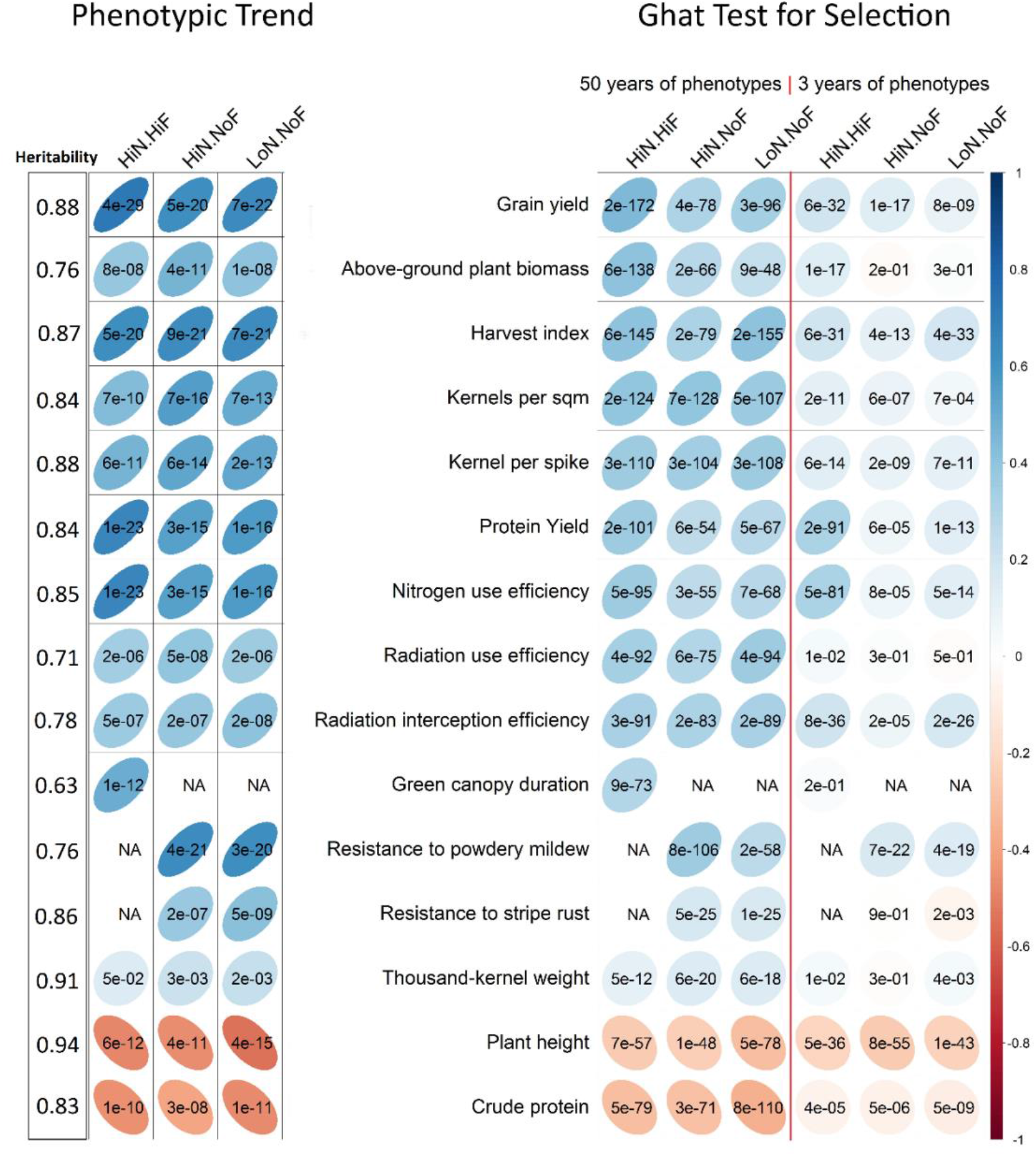
The power of Ghat test of selection in European winter wheat under three agrochemical treatments. 1: A high-intensity nitrogen supply along with best-practices fungicide, insecticide, and growth regulator applications (HiN/HiF), 2: A high level of nitrogen fertilization with a fungicide-free treatment (HiN/NoF), and 3: A low level of nitrogen fertilization with a fungicide-free treatment (LoN/NoF). **Left**: the phenotypic trend over the last 50 years. **Right**: The Ghat test of selection based on two analyses. “All Phenotypes” refers to estimating allelic effects using all available phenotypic information over the 50 years of wheat breeding; “Modern Phenotypes” refers to estimating allelic effects using modern phenotypic information only (2010 to 2013). Blue, positive selection; red, negative selection. Large light ellipses correspond to weaker selection intensities; thin dark ellipse correspond to more selection intensity. P-values printed on each cell on the left side of the figure (phenotypic trend section) correspond to the significance of the phenotypic trend. P-values printed on each cell on the right side of the figure (Ghat test section) correspond to implementing the Ghat permutation test.

#### Ghat test for selection

Fig 2 shows the results of the Ghat test when testing for selection between 1966 and 2013 in winter wheat breeding programs from mainly Western Europe. First, we investigated the precision and accuracy of the Ghat test by estimating the significance, and direction of selection and comparing the results to phenotypic progress over a time frame of 50 years. Overall, the selection size and direction on the 15 traits using all phenotypes were in the strong agreement with the phenotypic trend. Next, we evaluated the results from Ghat using phenotypic information from only the most-recent three years of the study (but all genotypic data were still included). Notably, this was also in a strong agreement with the phenotypic trend. Ghat successfully detected the signals of selection from the past in each trait, even when only modern phenotypes were utilized. For example, the results from the Ghat test indicated strong, directional selection for plant height and crude protein traits, which agrees with their phenotypic trends during the last 50 years[24]. Ghat showed very weak selection for thousand-kernel weight, which again aligned with this trait’s observed phenotypic trend [24]. The rest of the traits were under high to moderate directional selection. To validate the results of Ghat on wheat data, *spearman* correlations between the Ghat results from 3 and 50 years of phenotypes were estimated. A significant (r=0.57; P<0.001) correlation was found between Ghat values from 3 and 50 years of phenotypes, indicating that even in the absence of historical phenotypic information, Ghat can be used to understand historical patterns of selection.

#### Assessing population stratification

Our PCA showed little evidence for population stratification: no major groupings could be identified (Fig 3). Voss-Fels et al. (2019) [24] observed the same, and the authors explain that this observation may be caused by a large germplasm exchange between breeding companies. In the PCA, the first three principal components (PCs) explain together 16.05% of the variance, which do not correspond to their geographic location (Fig 3). This analysis was performed because excessive population stratification can cause erroneous signals of selection to be identified [16]. The minimal stratification observed in this PCA (Fig 3) and by Voss-Fels et al. (2019) [24] suggests this is not the case (Fig 3), which could be also explained by the strong exchange between the breeding programs [24].

**Fig 3.**
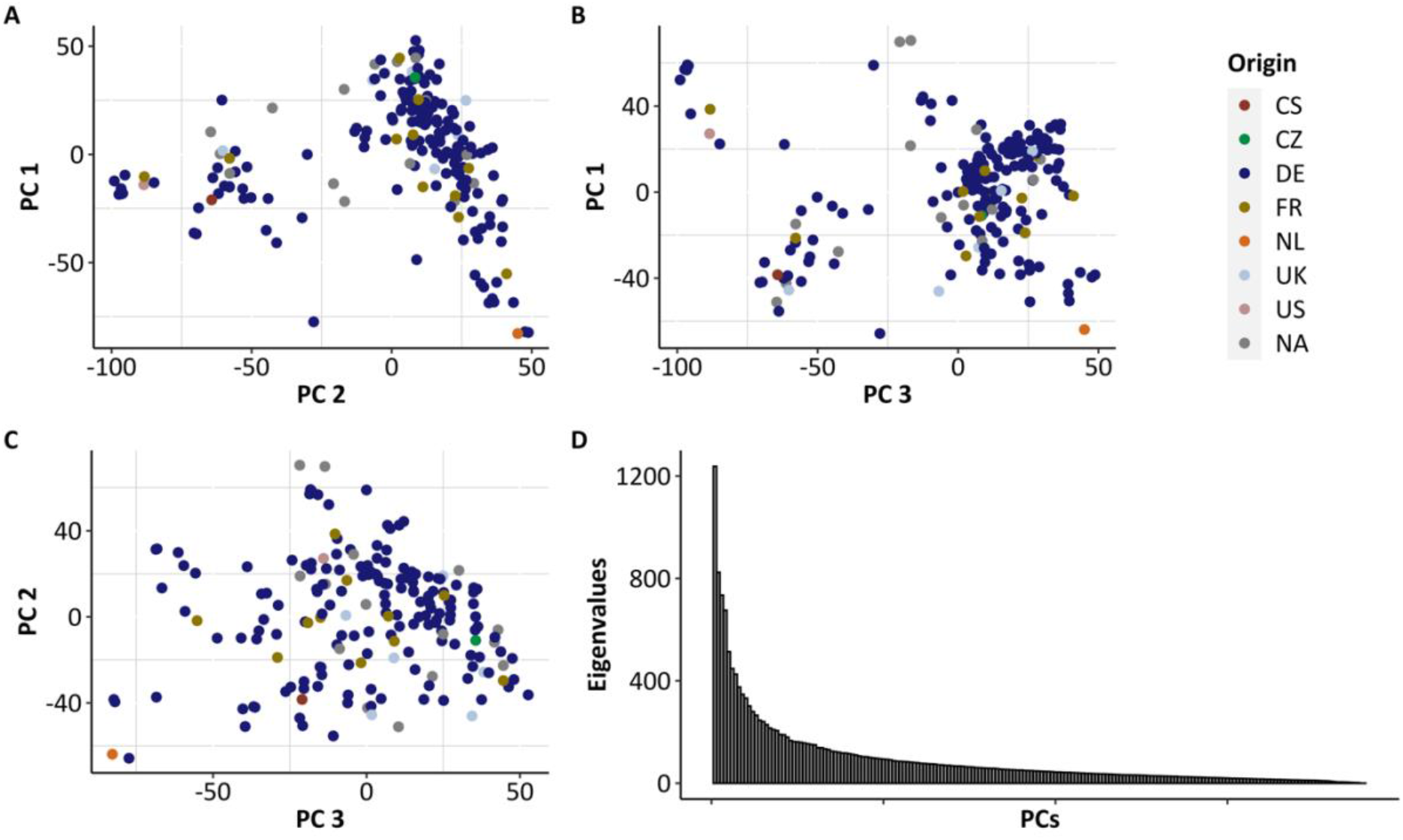
The results from the principal component analysis (PCA). A- PC1 plotted against PC2. B- PC1 plotted against PC3. C- PC2 plotted against PC3. D-The eigenvalues of all the detected PCs.

### Adaptation history of Western European winter wheat

To dissect the history of selection in wheat more precisely, we divided the 50 years of breeding into 10 five-year time periods and applied the Ghat test successively for each period. We excluded the green canopy duration trait from this analysis since it was only measured under one environmental condition. For the remaining 14 traits, we calculated the average phenotypic value across the three different environmental conditions. For example, during the period from 1966 to 1975, significantly selected traits according to the Ghat test included grain yield, harvest index (Fig 4-A), above-ground plant biomass, thousand-kernel weight (Fig 4-B), crude protein, nitrogen use efficiency (Fig 4-C), kernel per spike and resistance to stripe rust (Fig 4-D). However, during the same period, selection operated significantly in the negative direction against radiation use efficiency (Fig 4-A), thousand-kernel weight (Fig 4-B), plant height and resistance to powdery mildew (Fig 4-D). Therefore, by considering the single time periods of selection within the past 50 years (Fig 4), and regarding the traits which were selected within the different time periods, we can deduce information about the focus of the breeders during that time period or the environmental conditions during the time period. The number of cultivars available for the 10 adaptation periods are as follows: 18 cultivars from 1966 to 1975; 19 cultivars from 1976 to 1985; 25 cultivars from 1986 to 1995; 48 cultivars from 1996 to 2005; 81 cultivars from 2006 to 2015.

**Fig 4.**
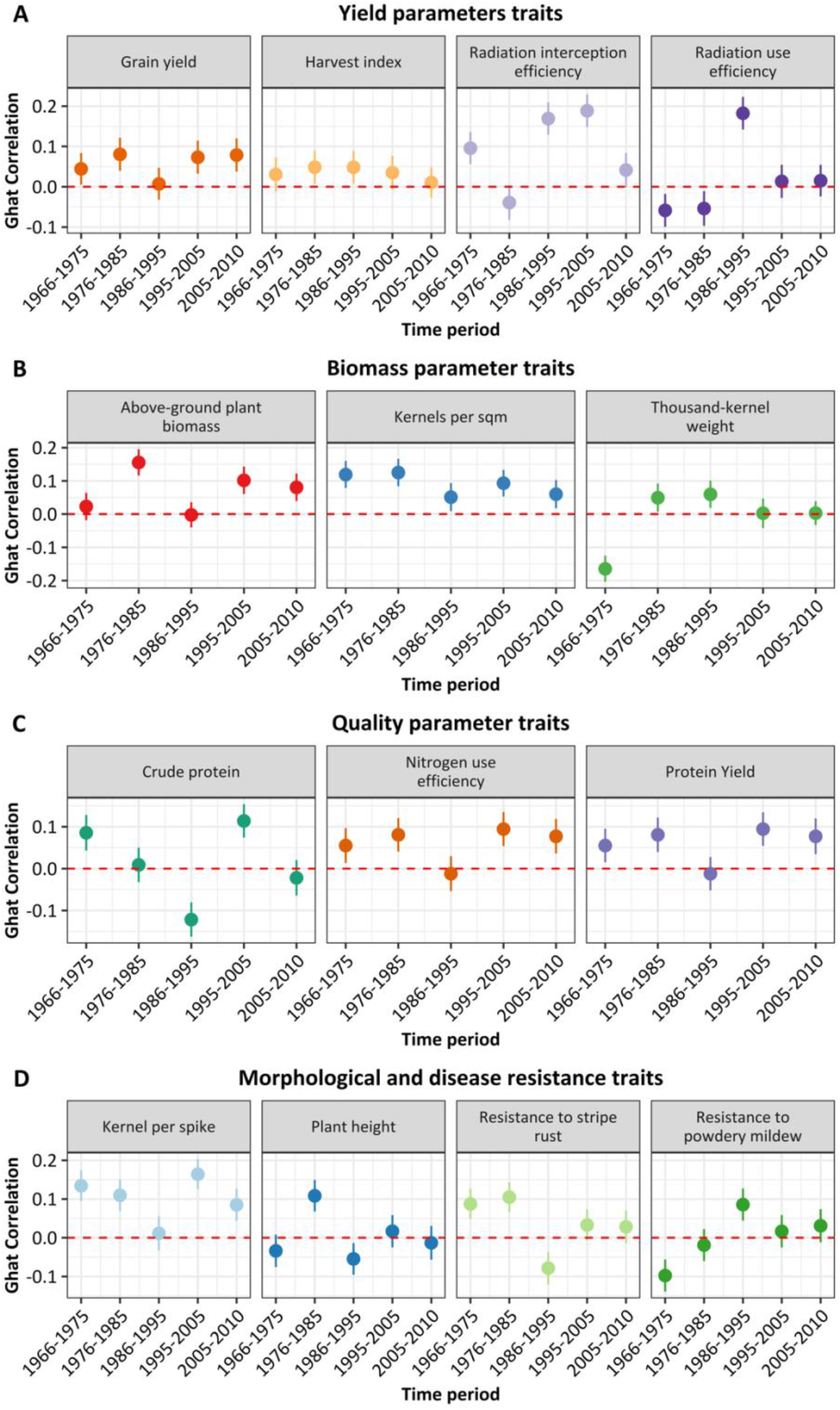
Ghat correlation estimates of the selection size and direction for 50 years of wheat breeding. Results are for 14 productivity traits in winter wheat from Western Europe measured during the past 50 years. Points above zero level (red dashed lines) indicate positive selection, while points under zero indicate negative selection; bold lines represent 99% confidence interval. A- Adaptation history for the **Yield parameters traits**. B- Adaptation history for the **Biomass parameters traits**. C- Adaptation history for **Quality parameters traits**. D- Adaptation history for **Morphological and disease resistance traits**.

## Part II: The power of Ghat to identify selection in a divergent of simulated dairy cattle population

To complement the above wheat analysis, we additionally generated a simulated cattle data set with divergent selection to further test and demonstrate the efficacy of Ghat. This allowed us to dissect the Ghat package and how it performs under different genetic architectures and experimental parameters, which are never perfectly known using real data. In this analysis, we assessed the impact of trait heritability (*h*^*2*^), sample size (*n*), trait polygenicity (number of QTL (*nQTL*)), and marker density (*MD*) on the performance of Ghat.

### Data Simulation

We simulated 30 bovine chromosomes of a length of 100 cM using QMsim [33]. Our simulations started with 100 historical generations of drift-only, and then we sampled two distinct populations to undergo selection. Population-A (unselected population) is a population that underwent complete random mating (i.e. no selection) for 20 generations; Population-B (selected population) is a population that underwent strong phenotypic selection for 20 generations (Fig 5). Aside from selection, Population-A and Population-B were identical and were treated identically during the simulated experiment. After simulating the populations, we applied Ghat to test for selection during the most recent 5 generations (generation #20 – generations #15) in both populations (unselected population and selected populations), creating a realistic situation in dairy cattle breeding. We estimated the allele effects using the genotypes of all 20 generations for more accurate estimation.

**Fig 5.**
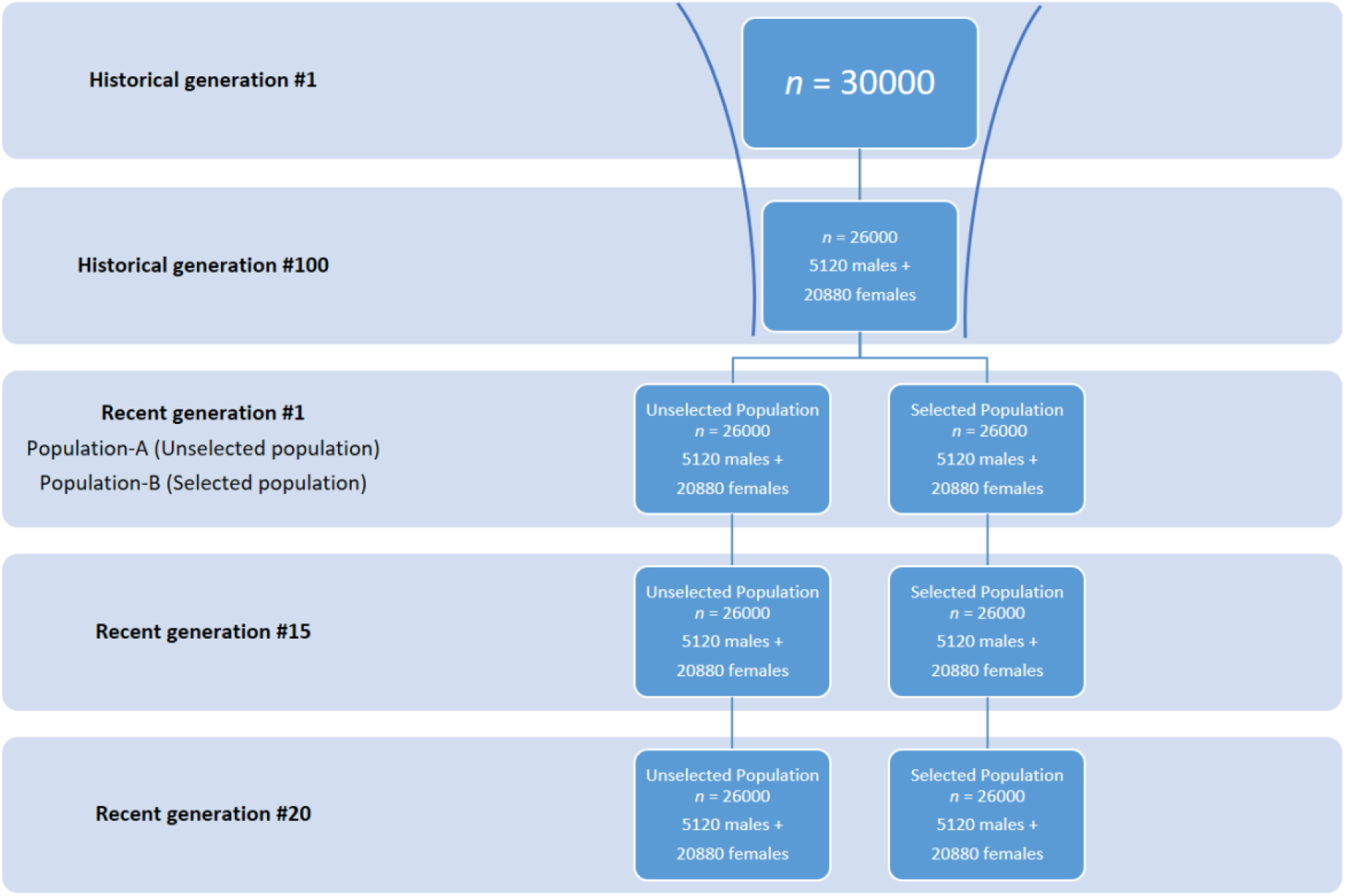
QMSim simulation scheme.

Four groups of datasets were simulated for studying the performance of Ghat (Table 2). Each group involved the creation of different datasets simulated with three constants parameters and one varying parameter (i.e. the parameter under investigation). Group 1 was simulated to investigate the effects of *n* in the Ghat test; we simulated ten datasets with a constant *h*^*2*^, *nQTL*, and *MD*, but with a different level of *n* (from 50 to 25600 animals) in each dataset. Then we measured the P values coming from using these different datasets with different *n*. Group 2 was simulated to investigate the effect of *h*^*2*^ on Ghat performance. For this, we simulated ten datasets with a constant *n, nQTL*, and *MD*, but with a different *h*^*2*^ (from 0.1 to 0.95) in each dataset. Group 3 investigated the effect of *nQTl* on Ghat performance. For group 3, we simulated ten datasets with a constant *n, h2*, and *MD*, but with a different level of nQTL (from 30 to 15360 QTL) in each dataset. Group 4 was simulated to investigate the effect of *MD* on Ghat performance. In this group we simulated ten datasets with a constant *n, h2*, and *nQTL* but with a different level of MD (from 30 to ∼ 1 million markers) in each dataset.

**Table 2:**
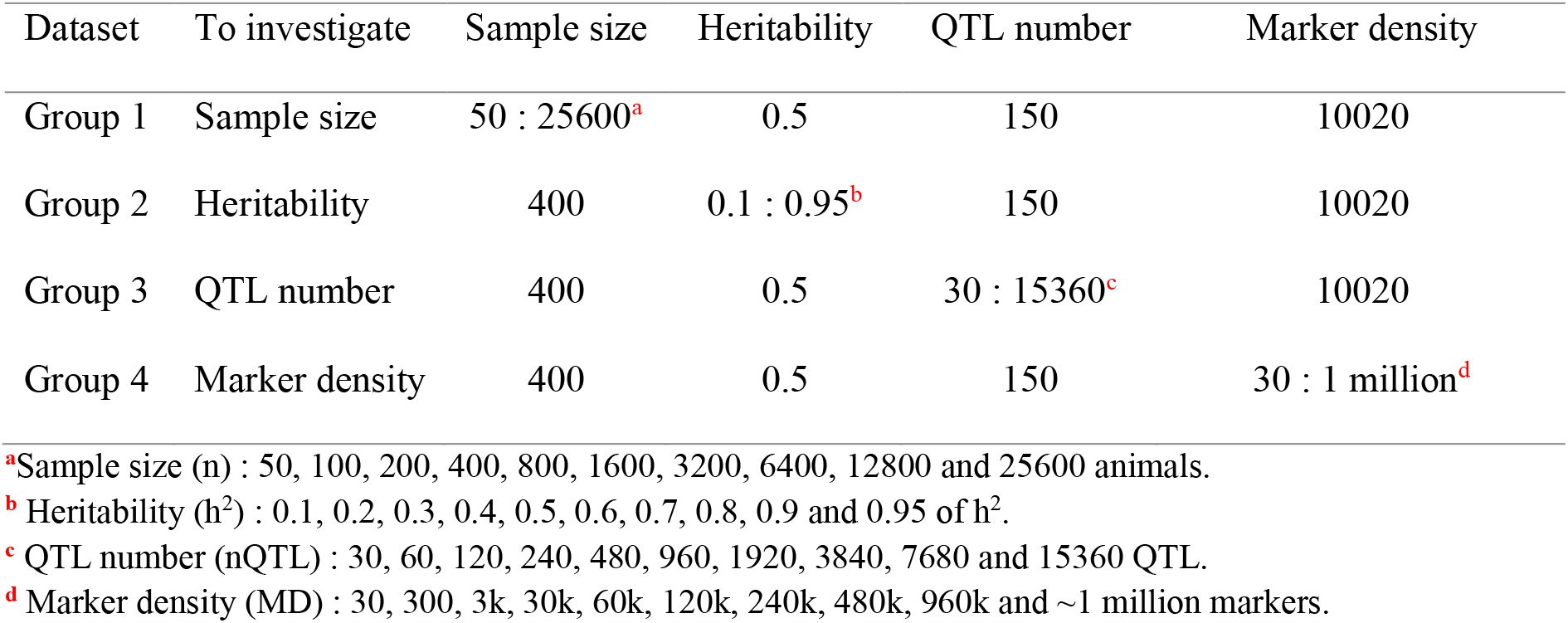
Presentation of all datasets used in the Ghat test.

| Dataset | To investigate | Sample size | Heritability | QTL number | Marker density |
| --- | --- | --- | --- | --- | --- |
| Group 1 | Sample size | 50 : 25600<sup>a</sup> | 0.5 | 150 | 10020 |
| Group 2 | Heritability | 400 | 0.1 : 0.95<sup>b</sup> | 150 | 10020 |
| Group 3 | QTL number | 400 | 0.5 | 30 : 15360<sup>c</sup> | 10020 |
| Group 4 | Marker density | 400 | 0.5 | 150 | 30 : 1 million<sup>d</sup> |

<sup>a</sup>Sample size ( $n$ ) : 50, 100, 200, 400, 800, 1600, 3200, 6400, 12800 and 25600 animals.
<sup>b</sup> Heritability ( $h^2$ ) : 0.1, 0.2, 0.3, 0.4, 0.5, 0.6, 0.7, 0.8, 0.9 and 0.95 of $h^2$ .
<sup>c</sup> QTL number ( $nQTL$ ) : 30, 60, 120, 240, 480, 960, 1920, 3840, 7680 and 15360 QTL.
<sup>d</sup> Marker density ( $MD$ ) : 30, 300, 3k, 30k, 60k, 120k, 240k, 480k, 960k and ~1 million markers.

### Method and implementation

Genomic prediction methods like BayesR, BayesB and BayesC assume that the allele substitution effects follow a distribution in which many effects are zero and some have moderate effects, subsequently these methods result in higher accuracy than the GBLUP/rrBLUP methods [34]. Since most of the bovine traits were simulated with a large number of QTL of small effect, allele substitution effects at every marker locus were estimated using BayesC [35]. The effective number of independent marker genome segments were estimated using SimpleM algorithm in R [23,36], and changes in allele frequencies were calculated with R [22,23]. All R-codes are publicity available in GitHub (https://github.com/Medhat86/Selection-and-adaptation-on-simulated-bovine-genome). Ghat is calculated as the summation of the estimated effect of every SNP scored multiplied by its effect size, according to model 1. We implemented identical analyses on selected and unselected populations across 1000 simulation replications for four different tests (*n, h*^*2*^, *nQTL and MD*) to evaluate the detection accuracy by evaluating true discovery rates and false positives. The confidence intervals were obtained from p-values of the same traits from the Ghat permutation test using the formula from Altman and Bland [37].

#### The effect of sample size on Ghat

Based on 1000 replications, Fig 6 shows the impact of *n* on the performance of Ghat through quantifying the P values of the Ghat test for selection (Fig 6). When we test the unselected population with a constant *h*^*2*^ = 0.5, *nQTL* = 150 and 10,020 markers, 99% of replicated traits were not significantly (p-value < 0.01) under selection. Conversely, in the selected population 99.3% of replicated traits found highly significant under selection (p-value < 0.01). The p-value also shows a decreasing trend with increasing *n* (from 50 to 25,600 animals) in the selected population. Moreover, the confidence interval decreases with increasing *n*.

**Fig 6.**
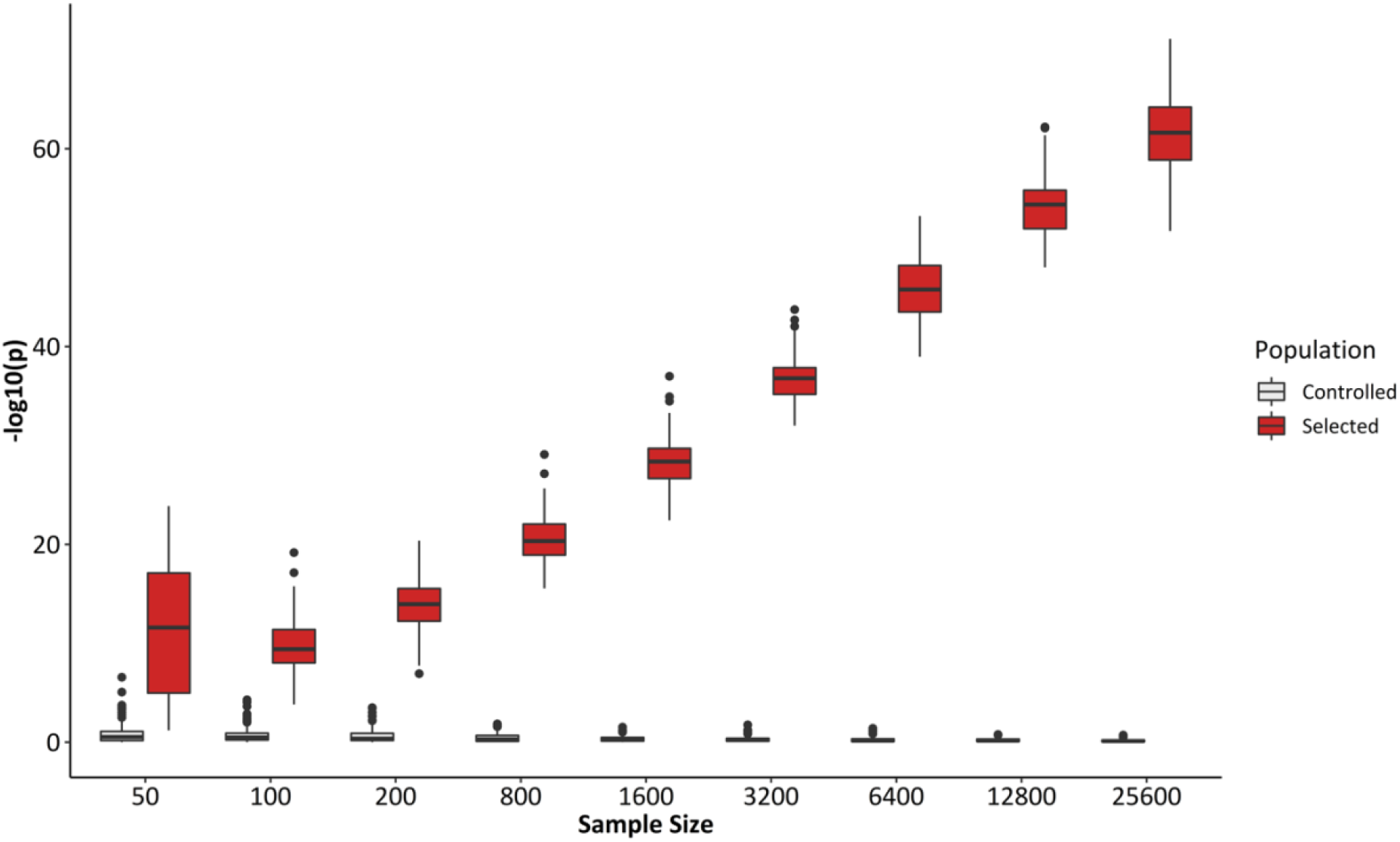
The effect of *n* on Ghat. Results are for 1000 replicated traits for testing the effect of *n* on Ghat-P.values, light blue for unselected population and dark blue for selected population.

#### The effect of heritability on Ghat

Based on 1000 replications, Fig 7 shows the impact of *h*^*2*^ on the power of Ghat (Fig 7). Higher *h*^*2*^ led to a more powerful test. When we tested the unselected population with a constant *n* = 400, *nQTL* = 150 and 10,020 markers, 100% of replicates were not significantly (p-value < 0.01) under selection. In the selected populations 100% of replicates were significant for selection (p-value < 0.01). The p-value shows a consistence decrease with increasing trait *h*^*2*^ (from 0.1 to 0.95 of *h*^*2*^).

**Fig 7.**
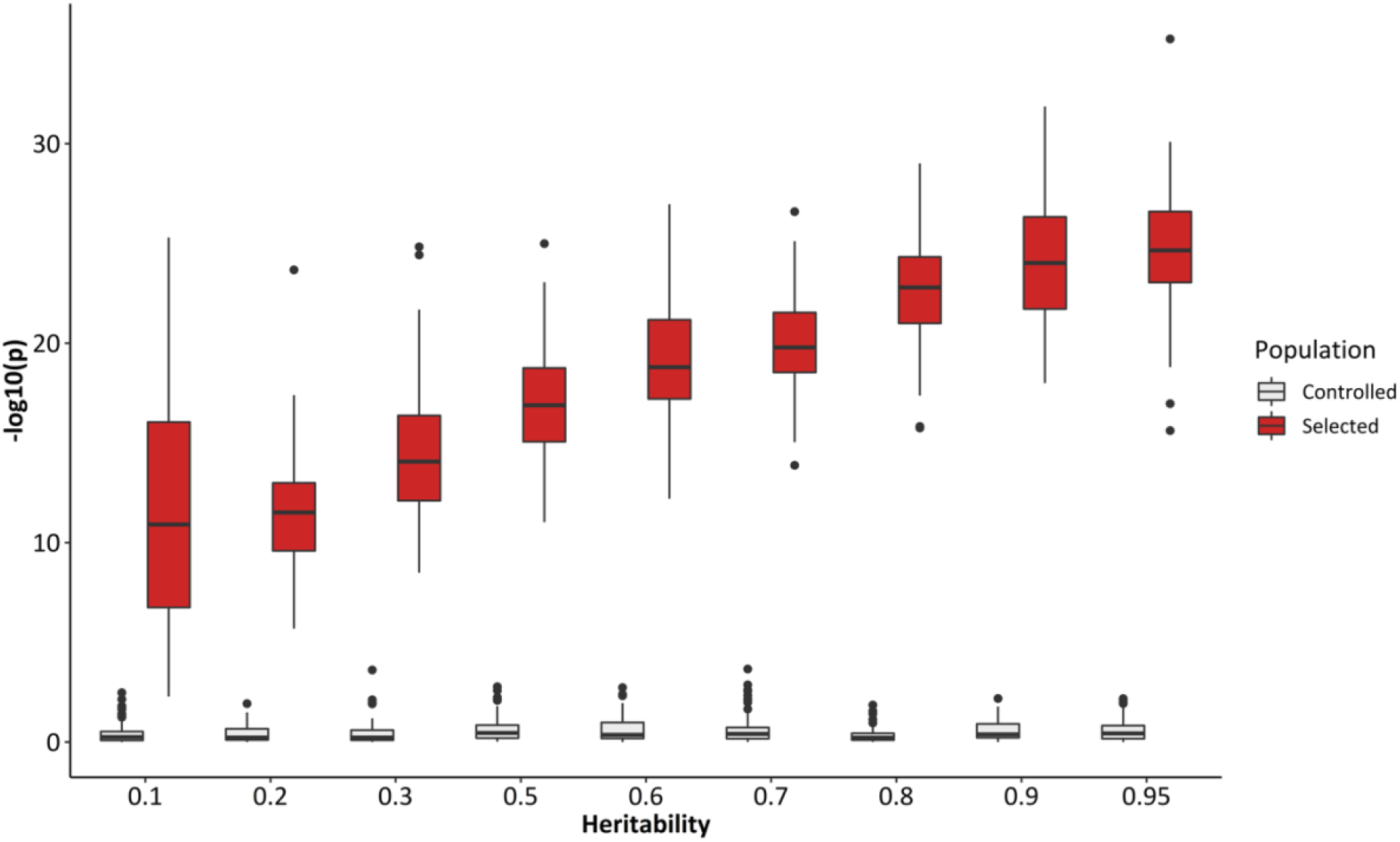
The effect of *h*^*2*^ on Ghat. Results depict the effect of *h*^*2*^ on the power of Ghat. Light green indicates the range of 1000 unselected population replicates and dark green indicates the range of 1000 selected population replicates.

### The effect of the number of QTL controlling a trait *nQTL* on Ghat

Based on 1000 replicates, Fig 8 shows the impact of *nQTL* on the power of Ghat. When we tested the unselected population with a constant *n* = 400, *h*^*2*^ = 0.5 and 10,020 markers, 99.7% of replicated traits were not significantly (p-value < 0.01) under selection. In selected population 100% of replicated traits found highly significant selection (p-value < 0.01). The power of the test increased with increasing the *nQTL* in selected population. This can be restated to emphasize that, all else being equal, Ghat is more able to detect selection on traits that are more complex (ie controlled by large number of QTL).

**Fig 8.**
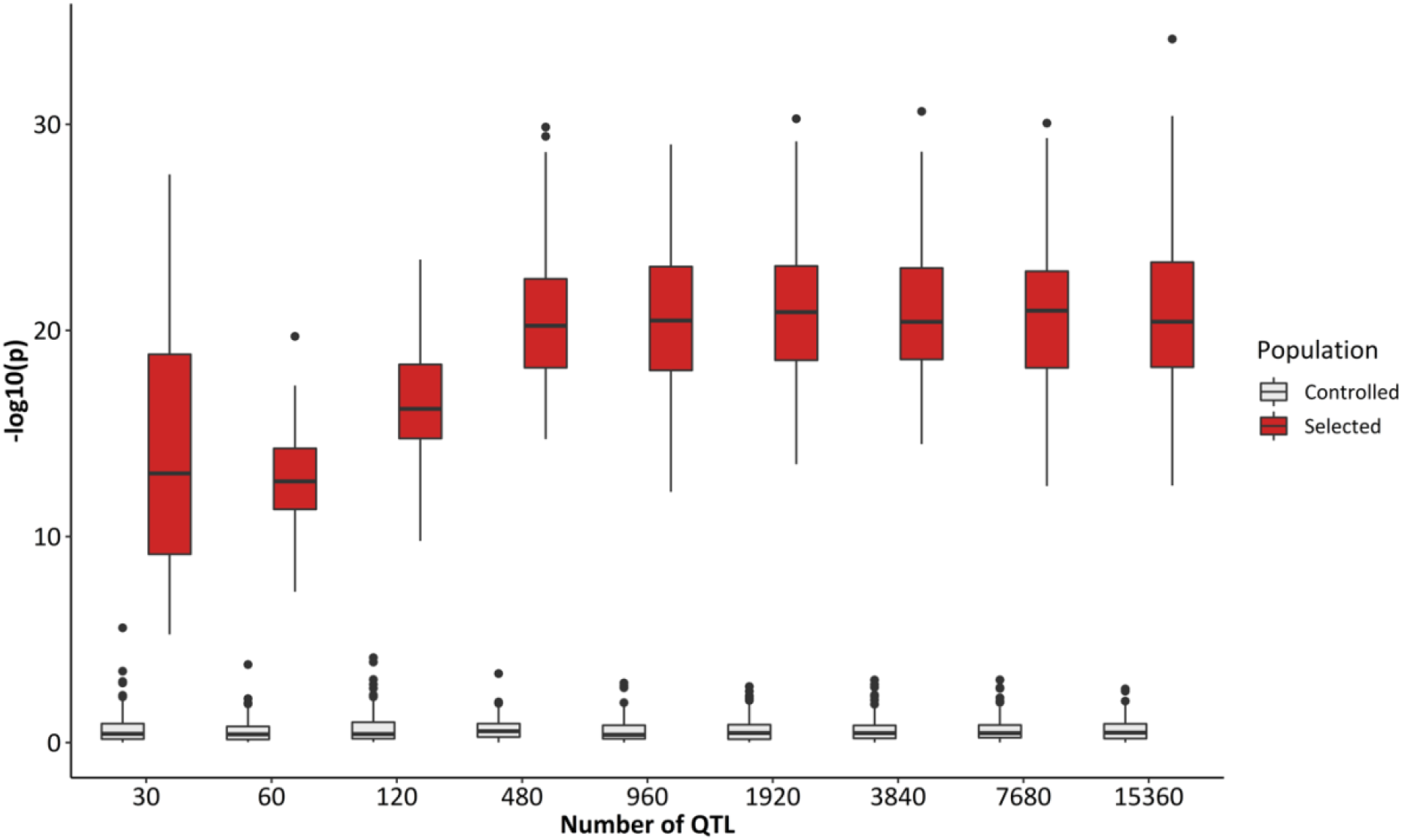
The effect of *nQTL* in the Ghat-test for selection. Results are for 1000 replicated traits for testing the effect of *nQTL* on Ghat-P.values, light red for unselected population and dark red for the selected population.

#### The effect of marker density on Ghat

Based on 1000 replications, Fig 9 shows the impact of *MD* on the performance of Ghat through quantifying the p-values of the Ghat-test for selection (Fig 9). When we tested the unselected population with a constant *n* = 400, *h*^*2*^ = 0.5 and 150 *QTL*, 90% of replicated traits were not significantly (p-value < 0.01) under selection. While, in selected population 100% of replicated traits were highly significant for selection (p-value < 0.01). The p-value also shows an inconsistent increase of significance along with an increasing MD in the selected population (from 30 to ∼ 1 million MD). This may be caused by multicollinearity in the data when there are a high number of markers (>120,000 MD) and relatively small n (400 animals).

**Fig 9.**
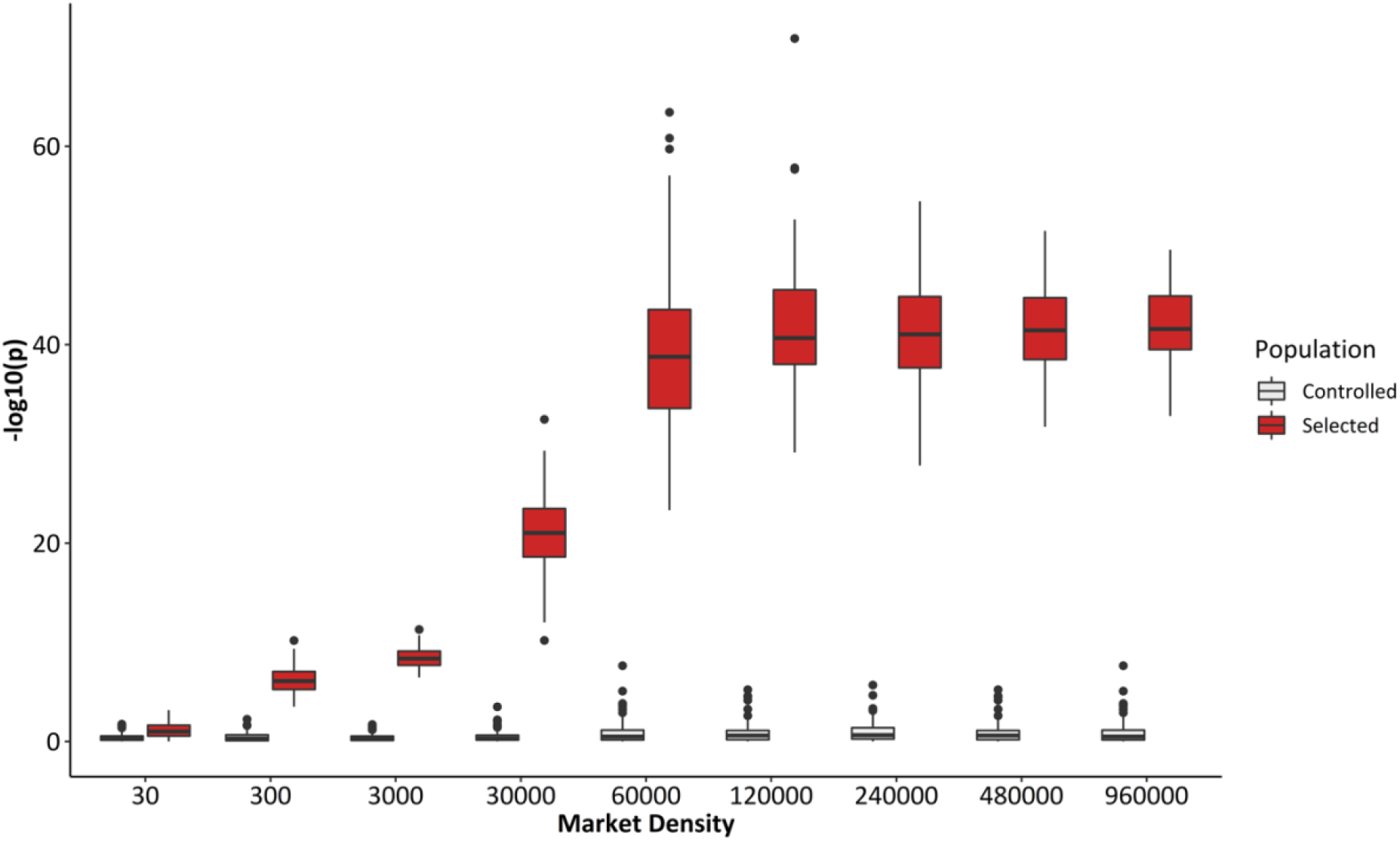
The effect of *MD* in Ghat-test for selection. Results are for 1000 replicated simulations testing the effect of *MD* on the power of Ghat. Gray indicates no-selection simulations and black indicates selection simulations.

#### Testing and system requirements

We have released three versions of Ghat package in CRAN (https://cran.r-project.org/package=Ghat) for the three major operating systems: MS Windows, Linux/Unix, and Mac OS. We have also released the source code on GitHub (https://github.com/timbeissinger/ComplexSelection) so that users can compile for specific platforms or modify the code as needed.

For users who have already computed allele effect and allele frequencies changes and are applying the Ghat test to these data, the computational cost of Ghat is unnoticeable. Practically, estimating Ghat for maize data [5] with 1000 permutations takes one second per one phenotype on a 2019 Dell XPS 23 with 2.4 Intel Core i7 processor.

### Future directions

As shown herein, the Ghat package can be powerfully used to detect selection on polygenic traits. Ghat identified polygenic selection on several candidate traits which were actually selected in winter wheat breeding programs. However, the wheat dataset included a relatively small number of markers (8,710) relative to current high-density marker panels and/or whole-genome sequencing approaches, so this analysis did not represent all the challenges associated with implementing Ghat in these settings.

To address this, we used a simulated SNP panel to further explore the utility of Ghat when high LD between markers is present. We found that Bayesian methods, such as BayesC [34,38], as implemented here, can be useful because of its ability to employ feature selection for marker number reduction [38–40]. BayesC does this by assuming that some fraction of the markers have zero effect on the trait of interest and forcing all other effects to equal zero. While Ghat proved to be powerful when paired with BayesC, Ghat is flexible and able to use marker effect estimates extracted from any prediction method. Further research should be conducted to determine the ideal feature-selection approaches to pair with Ghat.

Unlike methods for detecting selected loci, Ghat can quantify the direction of selection even in the absence of phenotypes from a pre-selection population. For breeders and breeding organizations, this represents a practical solution for tracing selection and adaptation history based on phenotypes measured from modern germplasm coupled with DNA samples saved from the past. From economical perspective, this may allow a retrospective evaluation of traits that were inadvertently selected in the past, and therefore may productively serve as the targets of intentional selection in the future. Advances in phenotyping technologies make measuring novel traits increasingly possible [41,42], and Ghat allows researchers to understand selection on traits that could not be measured previously for practical or technological reasons.

The Ghat test and implementation via the package described herein does not require immense sample size, individual genotypes and/or genome-wide significant SNPs, as demonstrated by our implementation with a relatively modest sample size of 191 wheat individuals genotyped for 8,710 SNPs. Notably, this approach is more powerful when traits are controlled by large number of QTLs [4]. These advantages allow the Ghat package to be used for identifying selection for various complex traits in other species of plants, animals, and humans.

## Data Availability

The Ghat R package described in this manuscript is open source and available on CRAN (https://cran.r-project.org/package=Ghat). All analysis and simulation scripts used in this manuscript are available on GitHub (https://github.com/timbeissinger/Maize-Teo-Scripts). The wheat dataset we utilized was previously published and released by [24].

## Acknowledgements

This project was supported with funds from the University of Göttingen, Faculty of Agriculture, the Center for Integrated Breeding Research, and the Division of Plant Breeding Methodology. We are indebted to the authors of [24] for making their data publicly available. Two anonymous reviewers provided suggestions that substantially improved this manuscript and we are grateful for their input. The authors gratefully acknowledge the hardware support from the High-Performance Computing team (HPC) at “Gesellschaft für wissenschaftliche Datenverarbeitung mbH Göttingen” (GWDG).

